# RP3Net: a deep learning model for predicting recombinant protein production in *Escherichia coli*

**DOI:** 10.1101/2025.05.13.652824

**Authors:** Evgeny Tankhilevich, Sergio Martinez Cuesta, Ian Barrett, Carolina Berg, Lovisa Holmberg Schiavone, Andrew R Leach

## Abstract

Recombinant protein expression can be a limiting step in the production of protein reagents for drug discovery and other biotechnology applications. We introduce RP3Net (Recombinant Protein Production Prediction Network), an AI model of small-scale heterologous soluble protein expression in *Escherichia coli*. RP3Net utilizes the most recent protein and genomic foundational models. A curated dataset of internal experimental results from AstraZeneca (AZ) and publicly available data from the Structural Genomics Consortium (SGC) was used for training, validation and testing of RP3Net. Set Transformer Pooling (STP) aggregation and Meta Label Correction (MLC) with large scale purification data enabled RP3Net to improve Area Under Receiver Operator Curve (AUROC) by 0.15, compared to the baseline model. When experimentally validated on an independent, manually selected set of 97 constructs, RP3Net outperformed currently available models, with an AUROC of 0.83, delivering accurate predictions in 77% of the cases, and correctly identifying successfully expressing constructs in 92% of cases.

## Introduction

### Motivation

The production of protein reagents is an essential part of the research and development process in the pharmaceutical and biotechnology industries. In drug discovery it is often a pre-requisite for screening and hit identification^1,2^. In living tissues, the target protein may occur in very small amounts alongside numerous other biomolecules. To be used for high-throughput screening of drug candidates, structural determination and the development of functional assays, the target protein needs to be expressed in a cell culture and purified. The ability to express a protein depends on multiple factors. First and foremost is the protein itself, but other factors include the cloning vector, the species and strain of the host cells, the codon optimisation algorithm, the use of tags and fusion proteins, and other experimental conditions ^3–21^. The choice of these parameters is often influenced by the details of the downstream experiments,^22^ making protein production time-consuming and error-prone, and often requiring multiple iterations and much trial and error. The purpose of this work is to develop a deep learning model to predict soluble protein expression in *E. coli* from the construct sequence, thus accelerating the timescales for protein production from months to weeks, cutting costs and reducing environmental impact.

A recombinant protein production experimental pipeline involves several steps^23^, including construct design, cloning, small-scale expression screening, progression of expressing constructs to large-scale purification and quality control (QC). Small-scale soluble expression screening, shown in **Fig. 1**, is crucial for assessing whether to progress the construct to large-scale production. First, cells are transfected with vectors (e.g. plasmids) carrying the cloned DNA of the protein of interest (step 1). Recombinant protein production is performed in deep well format (step 2). Cells are spun down and lysed (step 3). The lysate contains the total amount of protein produced. After an additional centrifugation step the soluble protein is found in the supernatant and the insoluble material is discarded in the pellet (step 3). Soluble protein is captured in a one-step purification using the histidine tag and immobilized metal affinity chromatography (IMAC^24^, step 4). The soluble protein yield and correct size are typically assessed by performing denaturing gel electrophoresis (SDS-PAGE) where yield and size are compared to a protein standard (step 5). The yield can be estimated by quantifying the amount of the target protein compared to the protein standard in the stained gel, using densitometric analysis. At this stage, it is important to record both positive (produced) and negative (failed to produce) experimental outcomes (step 6). Throughout the rest of this publication, unless stated explicitly, terms “protein production” and “protein expression” refer to this step of the experimental pipeline. Constructs that pass this small-scale screening are then typically progressed to large-scale purification and further downstream applications.

**Fig. 1.**
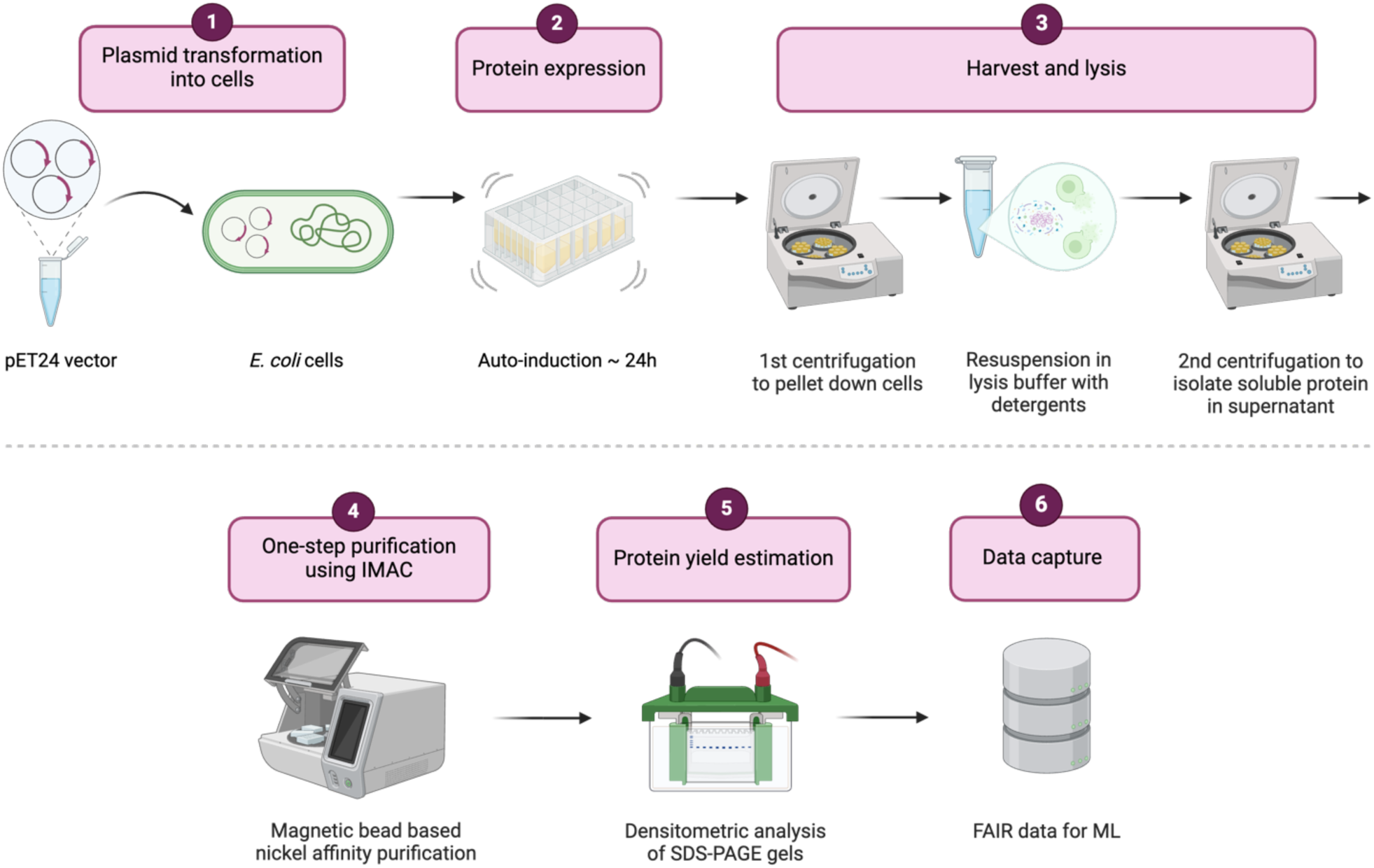
The experimental workflow for small-scale recombinant soluble protein production with one-step purification. After cloning, plasmids with the genetic material of the protein of interest are transfected into E. coli cells (step 1). The cells are grown for 24 hours (step 2). Harvesting involves two centrifugation stages: first to pellet down the cells, then, after lysis, to isolate the soluble protein in the supernatant (step 3). This is followed by IMAC purification (step 4), yield estimation via densitometric analysis of SDS-PAGE gels (step 5) and, finally, data capture for further analysis and machine learning (ML, step 6). Image generated with BioRender.com.

Protein and DNA foundation models (FMs) have become ubiquitous tools for predicting structural and functional protein properties from amino acid and/or nucleic acid sequences ^25–32^. These FMs are typically trained on large corpora of sequences, such as UniRef^33,34^, GenBank^35,36^, MGnify^37^ or BFD^38^ (Table 1). During training a portion of the sequence is masked out, i.e. each residue is replaced with a special “blank” character. The training objective then becomes to reconstruct the masked portion of the input, or “fill in the blanks”. This technique, referred to as language modelling task, originates from natural language processing^39^.

**Table 1.** Foundational models.

| Model | Type | Architecture | Training Data Sources | Training Data Size | Number of Parameters | Comments |
| --- | --- | --- | --- | --- | --- | --- |
| ESM2 650M<sup>26</sup> | Protein | Transformer | UniRef50, UniRef90 | 65M | 650M |  |
| ESM2 3B<sup>26</sup> | Protein | Transformer | UniRef50, UniRef90 | 65M | 2.8B |  |
| ESM3<sup>71</sup> | Protein | Transformer | Uniref, MGnify, JGI IMG/M, OAS | 2.78B | 1.8B | Licensing restrictions apply |
| ESMC 600<sup>72</sup> | Protein | Transformer | Uniref, MGnify, JGI IMG/M | 2.3B | 575M | Licensing restrictions apply |
| ESMC 300<sup>72</sup> | Protein | Transformer | Uniref, MGnify, JGI IMG/M | 2.3B | 333M | Licensing restrictions apply |
| ProtT5 XL<sup>27</sup> | Protein | Transformer | UniRef50 | 49M | 1.2B | Encoder only |
| ProtBert<sup>27</sup> | Protein | Transformer | UniRef100 | 217M | 420M |  |
| Protein-Bert<sup>28</sup> | Protein | Transformer | Uniref90 | 106M | 16M |  |
| HyenaDNA Medium<sup>29</sup> | DNA | Hyena | Human Genome hg38 | 3.2B bp | 24M | Maximum sequence length = 450K bases |
| HyenaDNA Large<sup>29</sup> | DNA | Hyena | Human Genome hg38 | 3.2B bp | 46M | Maximum sequence length = 1M bases |
| DNABert<sup>31</sup> | DNA | Transformer | Genomes from 136 species belonging to 6 classes | 32.49B bp | 117M |  |
| CaLM<sup>32</sup> | Codon | Transformer | European Nucleotide Archive, coding sequences | 8.7M | 85M |  |

ESM^25,26,40^, ProtBert^27^ and ProteinBert^28^ are examples of protein FMs; DNABert^30,31^ is a popular DNA FM. These models are all based on Transformer deep learning architecture^41^ with different number of layers, feature dimensions and other details. HyenaDna^29^ is another DNA FM that uses a different architecture.

The intermediate layers of foundational models yield a residue-level sequence representation that can be used to predict the protein property of interest, such as secondary or tertiary structure, binding affinity, fluorescence, thermodynamic stability, solubility, etc^25,26,38,40,42–47^. The experimental datasets that describe these properties typically contain orders of magnitude fewer entries when compared to the sequence corpora. This scarcity of experimental datasets often makes it unfeasible to train large foundational models from scratch for predicting protein properties. Such models are usually pre-trained with the language modelling objective on the large corpora first, and then further trained to predict the property of interest using the smaller dataset^42,43,47^. This final training step is referred to as fine-tuning. RP3Net follows this architectural blueprint by encoding the biological sequence with a foundational model, feeding this encoding through an aggregation layer to obtain a global representation for the entire construct, and then applying a fully connected classification head to compute the predicted probability of recombinant expression in *E. coli* (**Fig. 2A**) as a binary outcome.

**Fig. 2.**
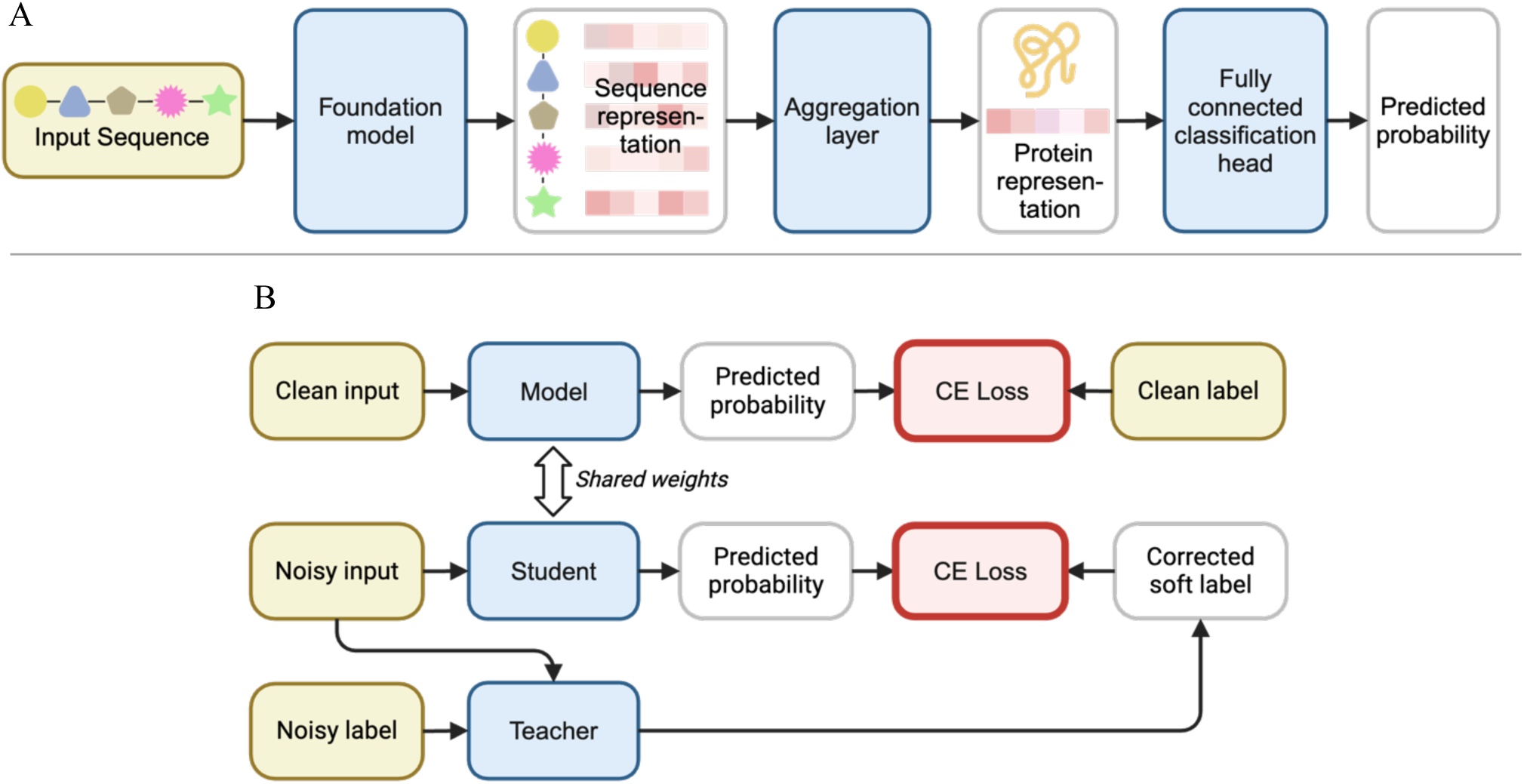
**A**. Architecture diagram of RP3Net. The input biological sequence is encoded by the foundation model to obtain a sequence representation, where each residue/codon/nucleotide is represented by a vector. The aggregation layer builds a global protein representation vector from the sequence representation. The predicted probability of successful recombinant expression of the protein in E. coli is computed by the fully connected classification head from the protein representation. **B.** Training with meta label correction on a mixture of clean and noisy data. The standard training setup, where the model loss on clean inputs and labels is minimised with gradient descent, is shown in the top row. A special “teacher” model is trained to predict the corrected labels from the noisy input and labels. These corrected labels, along with noisy inputs, serve as inputs for training the “student” model. The latter model has the same architecture and weights as the “clean” model. The bi-level optimisation algorithm that makes sure that the corrected labels do not deviate from the (unknown) clean labels, relies on using the cross entropy (CE) loss. Model components with trainable weights are shown as blue boxes. Training data is shown as yellow boxes. Images generated with BioRender.com.

Although, in theory, a soluble protein production fine-tuning dataset could be designed and experimentally generated from scratch, in practice it would be too time-consuming and expensive. Moreover, there already exist publicly available datasets of protein expression that contain the results of experiments worth millions of dollars and representing years of lab work^48–50^. For training and evaluating RP3Net, the internal AstraZeneca (AZ) small-scale expression screen data is combined with datasets from the Structural Genomics Consortium (SGC), specifically their sites in Stockholm^4,51^ and Toronto^23,50^. The experimental pipeline for generating data from AZ and SGC Stockholm has been already discussed above, see **Fig. 1**, step 6. SGC Toronto captures the results of large-scale protein purification.

### Existing work

A number of models that predict soluble expression from construct sequence have been published in recent years. Most of these systems use datasets derived from the Protein Structure Initiative (PSI) compendium, also referred to as TargetTrack ^48,49^. PSI was an experimental research effort run across multiple laboratories in 2000-2017, with the objective of determining protein structures and depositing them in the Protein Data Bank (PDB)^52,53^. This dataset records the pipeline position, i.e. the experimental stage where the work was terminated, for each target and construct. For example, if a construct was selected and cloned, but could not be expressed, its pipeline position would be recorded as “cloned”. For another construct that has been selected, cloned, expressed and purified, but could not be crystallised, the pipeline position would be “purified”, etc. One limitation of using TargetTrack data in this work is that in this dataset a genuine inability to express the construct under given experimental conditions can be confused with stopping to pursue the construct for other reasons (for example there being another well-behaving construct for the same target). Different labs that have provided data for TargetTack were using different experimental pipelines: sometimes small-scale expression screening as shown in **Fig. 1**, but also large-scale purification results as SGC Toronto, or the results of running SDS-PAGE on unpurified cell lysate.

It is important to make a distinction between solubility as a general physical property of the protein, which can be measured, for example, as peak concentration in the solution, and the ability to achieve soluble expression of the protein under given experimental conditions, which is a binary outcome that is modelled in this work. A protein that is generally soluble could still fail to express, for example because it is toxic for the host cells, or because the chaperones that are required for forming the correct structure are missing, or due to other reasons. Solubility is thus a necessary but insufficient condition for soluble recombinant protein production.

NetSolP^44^ uses PSI data to evaluate multiple transformer-based models available at the time for predicting soluble expression. NetSolP outputs two scores: solubility and “usability”, the latter being a combined predictor of solubility and the ability of a protein to be expressed.

PLMC^46^ and SADeepCry^45^ also use data derived from TargetTrack and a Transformer architecture but output the pipeline position given the construct sequence. PPCPred^54^, PredPPCrys^55^, Crysalis^56^ DCFCrystal^57^ are examples of older, simpler models that predict pipeline position, trained on various subsets of TargetTrack. SoluProt^58^ uses a different PSI-based dataset with a Gradient Boosted Machine (GBM) model^59^ and global features based on relative amino acid frequencies, predicted physicochemical properties, similarity to *E. coli* proteome and output of various other bioinformatics tools to predict soluble expression.

CamSol^60,61^ is a well-established relative solubility prediction tool for libraries of similar protein sequences. There are many other solubility predictions that use deep neural networks, such as GPSFun^62^ and PLM_Sol^63^. A few methods exist for modelling expression and solubility of human antibodies, but their experimental protocols differ substantially from *E. coli*-based expression analysed in this work^64,65^.

## Results and discussion

### RP3Net with fixed foundation model weights outperforms decision trees with global protein features

The RP3Net architecture with fixed foundation model weights and mean pooling (**Fig. 2A**) was used for selecting the best performing foundation model. This architecture is denoted as Model A (**Table 2**). The models were trained and evaluated on SGC Stockholm dataset, with five-fold cross validation. SGC Stockholm was used because this dataset is of medium size, compared to AZ, which is much smaller, and SGC Toronto, which is much larger. This data set is also the only one of the three that provides DNA sequences for all the constructs.

**Table 2.** RP3Net training and architecture configurations.

| **RP3Net Model** | **Aggregation** | **FM weights** | **Meta Label Correction** |
| --- | --- | --- | --- |
| A | Mean | Frozen | No |
| B | STP | Frozen | No |
| C | STP | Fine-tuned, LoRA | No |
| D | STP | Fine-tuned, LoRA | Yes |

A gradient boosted decision tree (XGBoost^59^) with global protein features as inputs was used as a baseline model. As shown in **Fig. 3**, Model A with any protein FM outperforms the baseline model. Out of all the tested DNA and codon FMs, only CaLM shows better results than the baseline. A plausible explanation for this observation is due to the datasets used for pre-training the FMs: CaLM was pre-trained on coding sequences from ENA, whereas other DNA FMs were pre-trained on a mixture of coding and non-coding sequences.

**Fig. 3.**
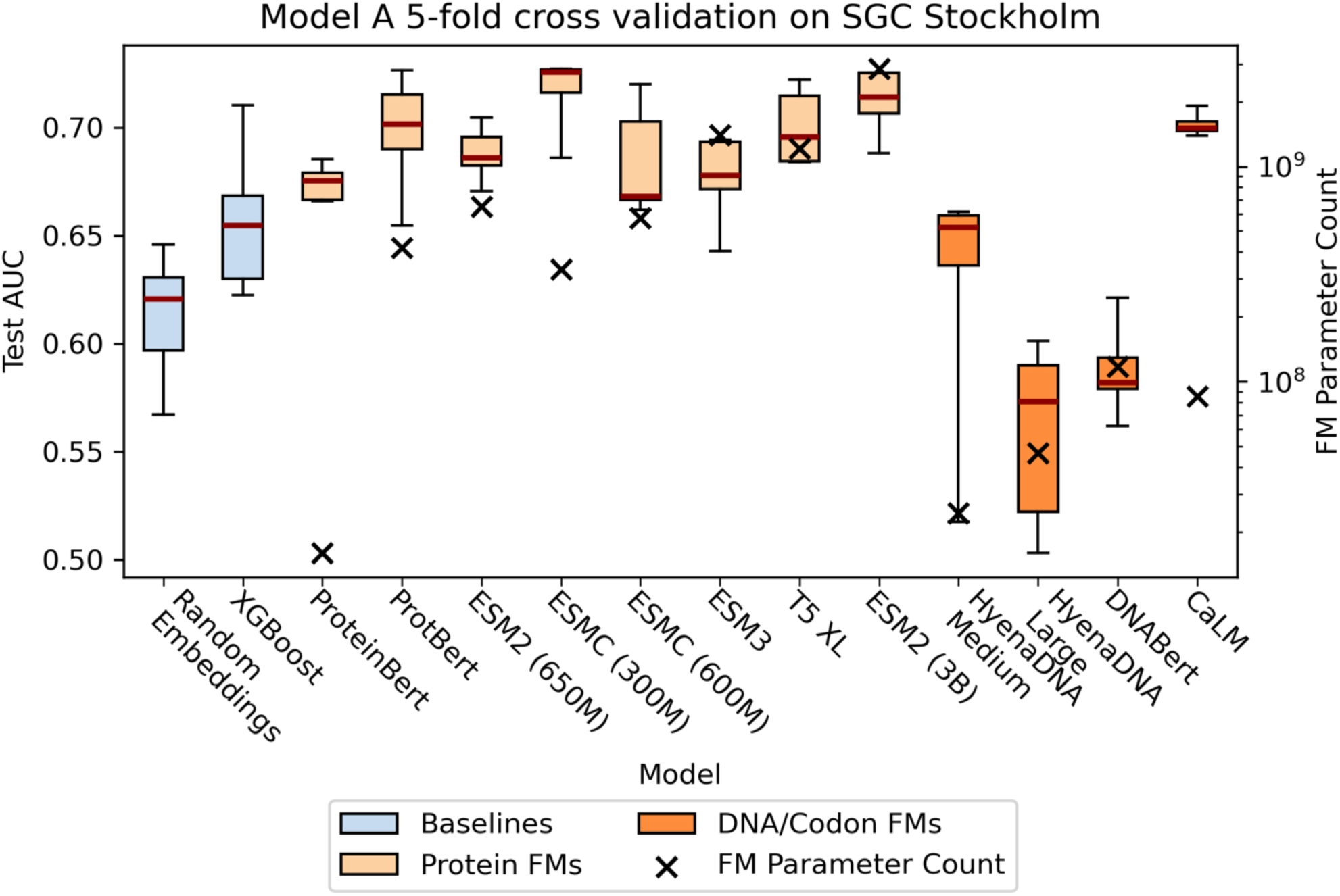
Performance of RP3Net with fixed foundation model (FM) weights and mean pooling (Model A) on SGC Stockholm, along with FM parameter count. On the left y-axis each boxplot shows the area under Receiver Operator Curve (AUROC) of Model A with a particular FM, evaluated on SGC Stockholm test data, with five-fold cross validation. On the right y-axis, black squares show the number of trainable parameters of the FM, in log scale. ESM2 (650M) and CaLM were selected for further analysis based on performance, consistency, parameter count and licensing restrictions. “Random embeddings” means using random residue embeddings instead of a foundation model. The source data for all charts is available in the supplement.

RP3Net performance also varies depending on the training data subset that was used, sometimes dramatically. For example, for the more consistent FMs, such as ESM2 (650M) and CaLM, the difference between the best and the worst runs is 0.03 and 0.01, respectively, whereas for HyenaDNA Medium the difference is 0.14.

The number of trainable foundation model parameters is used to indicate the resource requirements for fine-tuning the FM (compute time, memory). The foundation model for subsequent evaluation was chosen based on the pragmatic trade-off between performance, training complexity and licensing constraints (see **Table 1**). We selected ESM2 with 650 million parameters. The simple Model A training protocol, applied to this FM, achieves an average increase in AUROC of 0.03, compared to the baseline model.

### Performance on different data sources reveals dependency on dataset size

Model A performance on SGC Stockholm dataset can be improved by replacing the mean pooling aggregation layer with a more sophisticated set transformer pooling (STP^66,67^). This configuration is denoted as Model B. The main difference between mean pooling and STP is that, whereas the former just takes an average across the sequence, giving each residue the same weight, STP uses context-dependent weights for residue representations, by computing multiheaded attention (MHA^41^, see Methods) between a special parameter, called the seed vector, and the output of the foundation model. The seed vector is updated during training with gradient descent, along with the rest of the model parameters.

Model B gives an AUROC improvement of 0.01% over Model A when trained and evaluated on SGC Stockholm (**Table 3**). The performance of Model B on other data sources varies with an AU-ROC of 0.59 on the AZ dataset and of 0.84 on SGC Toronto. A plausible reason for this variation is dataset size. Training Model B on the combined AZ and SGC Stockholm data improves evaluation of AZ to 0.73, which is almost the same as evaluating the same model on SGC Stockholm. Adding SGC Toronto to the training data does not improve the evaluation results significantly for any data source.

**Table 3.**
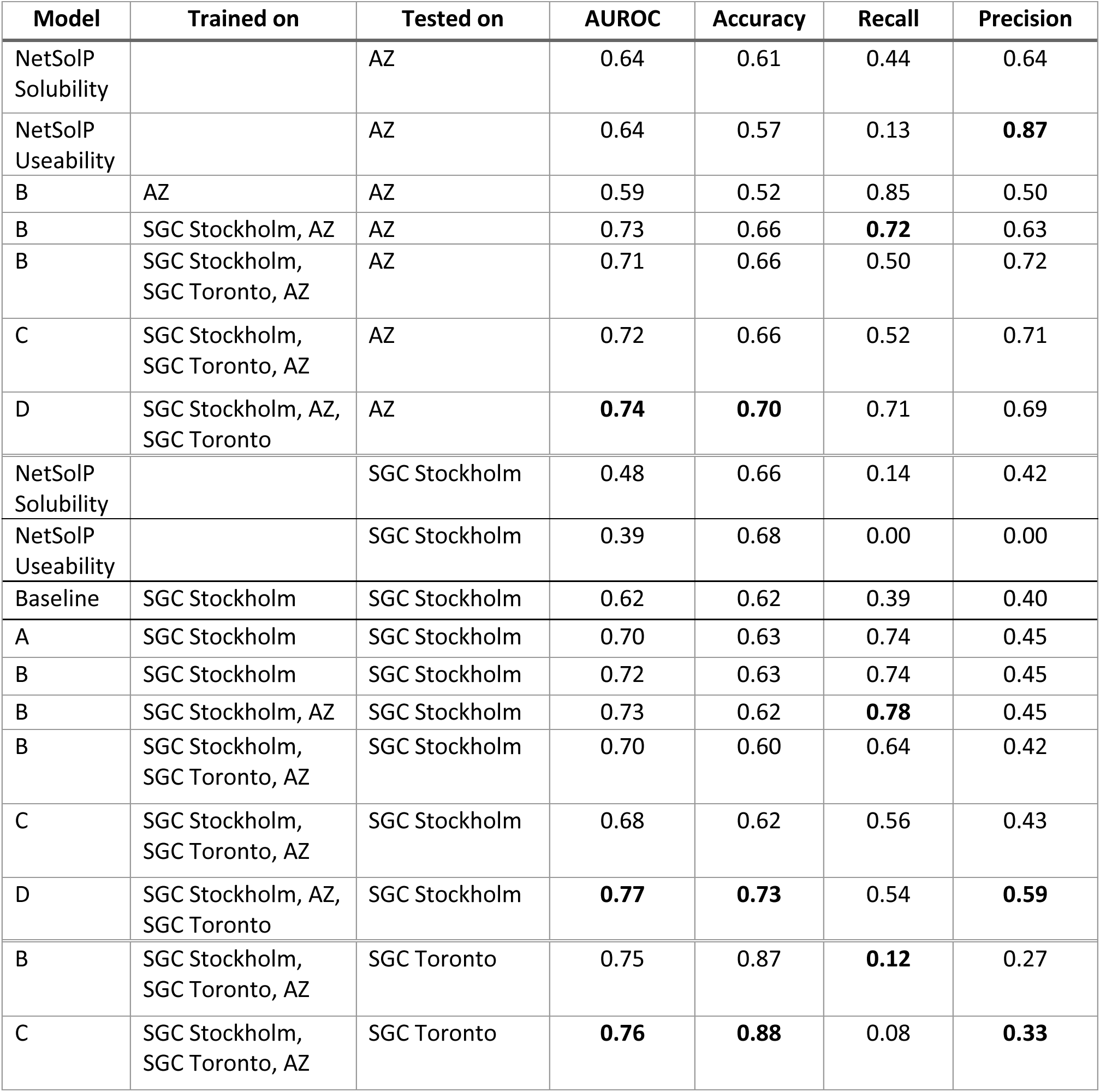
Results of evaluating different RP3Net models trained on different data sources, versus the baseline model and third party predictors.

| Model | Trained on | Tested on | AUROC | Accuracy | Recall | Precision |
| --- | --- | --- | --- | --- | --- | --- |
| NetSolP Solubility |  | AZ | 0.64 | 0.61 | 0.44 | 0.64 |
| NetSolP Useability |  | AZ | 0.64 | 0.57 | 0.13 | **0.87** |
| B | AZ | AZ | 0.59 | 0.52 | 0.85 | 0.50 |
| B | SGC Stockholm, AZ | AZ | 0.73 | 0.66 | **0.72** | 0.63 |
| B | SGC Stockholm, SGC Toronto, AZ | AZ | 0.71 | 0.66 | 0.50 | 0.72 |
| C | SGC Stockholm, SGC Toronto, AZ | AZ | 0.72 | 0.66 | 0.52 | 0.71 |
| D | SGC Stockholm, AZ, SGC Toronto | AZ | **0.74** | **0.70** | 0.71 | 0.69 |
| NetSolP Solubility |  | SGC Stockholm | 0.48 | 0.66 | 0.14 | 0.42 |
| NetSolP Useability |  | SGC Stockholm | 0.39 | 0.68 | 0.00 | 0.00 |
| Baseline | SGC Stockholm | SGC Stockholm | 0.62 | 0.62 | 0.39 | 0.40 |
| A | SGC Stockholm | SGC Stockholm | 0.70 | 0.63 | 0.74 | 0.45 |
| B | SGC Stockholm | SGC Stockholm | 0.72 | 0.63 | 0.74 | 0.45 |
| B | SGC Stockholm, AZ | SGC Stockholm | 0.73 | 0.62 | **0.78** | 0.45 |
| B | SGC Stockholm, SGC Toronto, AZ | SGC Stockholm | 0.70 | 0.60 | 0.64 | 0.42 |
| C | SGC Stockholm, SGC Toronto, AZ | SGC Stockholm | 0.68 | 0.62 | 0.56 | 0.43 |
| D | SGC Stockholm, AZ, SGC Toronto | SGC Stockholm | **0.77** | **0.73** | 0.54 | **0.59** |
| B | SGC Stockholm, SGC Toronto, AZ | SGC Toronto | 0.75 | 0.87 | **0.12** | 0.27 |
| C | SGC Stockholm, SGC Toronto, AZ | SGC Toronto | **0.76** | **0.88** | 0.08 | **0.33** |

### Meta label correction with purification data yields a 0.04 increase in AUROC on SGC Stockholm

Both Model A and Model B are fine-tuned on soluble protein expression data with frozen parameters of the foundation model. Unfreezing these parameters (Model C) and training on the full dataset leads to overfitting: perfect performance is quickly achieved on the training data set (AUROC≍1.0), but on the validation and test sets the AUROC remains below 0.75. Training Model C on individual data sources also leads to overfitting, as expected.

This could be explained by the fact that the datasets contain the results of slightly different experiments. The SGC Toronto dataset reports results of large-scale purification, whereas both AZ and SGC Stockholm report small-scale expression testing captured with one-step purification. Although the exact experimental conditions, materials and methods used for purifications were not available during model development, it is safe to assume that the SGC Toronto conditions are quite different from the small-scale expression testing. A natural question arises: given the construct sequence from SGC Toronto, and its binary purification result, what would be the result of small-scale expression testing this construct under the conditions of SGC Stockholm or AZ?

We address this within the Meta Label Correction framework (MLC^68,69^), where a large, noisy data set is used to aid training the model on a small, clean set. Rather than adding the noisy data directly to the training set, a special model is trained to predict the corrected label from the noisy input and noisy label. This is referred to as the “teacher model”. The corrected labels are used to train the “student model”, along with clean inputs and clean labels (**Fig. 2B**). In our setup, SGC Toronto large scale purification data set is used to train the teacher model, and a union of SGC Stockholm and AZ small-scale expression data is used to train the student model.

Using MLC with large scale purification data (Model D) achieves in AUROC of 0.74 on AZ dataset. This is an improvement of 0.01 compared to the second-best result of Model B on AZ data. When evaluating Model D on SGC Stockholm, AUROC reaches 0.77, which is an improvement of 0.04 over the next-best result. We have also observed that the MLC model is more robust across different sequence clusters – training, validation and testing – than other models, which tend to overfit the training data. The MLC framework thus allows utilising large scale purification data to improve modelling of small-scale expression testing, whereas simple transfer learning (Model B or Model C trained on all sources) fails to achieve that outcome.

### Prospective experimental validation of the model shows AUROC of 0.83

To establish the utility of RP3Net for drug discovery projects, in addition to the normal train-validate-test model development loop, we have conducted prospective model evaluation in a real-life scenario. A set of 46 proteins was curated from the human proteome to include viable drug targets, whilst avoiding proteins with prior published evidence of successful expression. We started by generating two full length constructs per target (with a 6-His affinity tag placed at the N- or C-Terminal) and running RP3Net on them. If both constructs were predicted not to express, we generated trimmed constructs, ran these through the model, and, if they were predicted to express, included them in the dataset (see Methods). This resulted in a total of ninety-seven constructs for the experimental validation dataset, eight of which were generated by the trimming process (see Methods). The constructs were cloned and expressed in *E. coli* at the AZ protein production facility. 54% of the constructs passed small-scale expression screening, including one-step purification by affinity chromatography. The remaining 46% were annotated as either “Ambiguous” or “Not Passed”.

The performance of RP3Net models B and D was compared with the baseline model, and two third-party predictors: SoluProt^58^ and NetSolP^44^ (**Table 4**). The highest AUROC of 0.83 is achieved by RP3Net model D. This is 0.06 better than the next-best result (RP3Net B trained on AZ and SGC Stockholm), and 0.08 better than the best third-party predictor (NetSolP useability). RP3Net D showed an accuracy of 0.77 when the score cut-off of 0.5 was used, and accuracy of 0.81 with the cut-off set to 0.79.

**Table 4.** Results of experimental validation of RP3Net, the baseline model, and third-party predictors.

| Model | Trained on | AUROC | Accuracy | Recall | Precision |
| --- | --- | --- | --- | --- | --- |
| Soluprot |  | 0.64 | 0.57 | 0.83 | 0.57 |
| NetSolp Solubility |  | 0.66 | 0.60 | 0.79 | 0.59 |
| NetSolp Useability |  | 0.75 | 0.57 | 0.23 | **0.86** |
| Baseline | SGC Stockholm | 0.67 | 0.65 | 0.58 | 0.71 |
| RP3Net B | AZ | 0.69 | 0.65 | 0.88 | 0.62 |
| RP3Net B | AS, SGC Stockholm, | 0.77 | 0.71 | **0.94** | 0.66 |
| RP3Net B | AZ, SGC Stockholm,<br>SGC Toronto | 0.76 | 0.69 | 0.65 | 0.74 |
| RP3Net D | AZ, SGC Stockholm,<br>SGC Toronto MLC | **0.83** | **0.77** | 0.92 | 0.73 |

For the subset of eight trimmed constructs, RP3Net D shows an accuracy of 0.5 with score cut-off of 0.5, and accuracy of 0.62 with score cut-off of 0.79. This could be an artefact of the small evaluation set, or that RP3Net does not consider if sequences will fold into stable protein domains. Curiously, the trimmed constructs that did result in soluble protein also contained degradation products (supplementary figure 1). With the score cut-off of 0.5 the model predicts all trimmed constructs to express, whereas in fact only four out of eight were expressed successfully.

Performance on trimmed constructs could thus be considered an area for improvement. However, considering the small number of trimmed constructs, and the model accuracy (0.77) and precision (0.73) on the larger experimental validation set, it could be argued that an experimental scientist would still find the modelling results helpful. An “overconfident”, high recall, model that predicts too many positives, which are then partly confirmed in the laboratory, is preferrable to a model that misses out constructs that would have expressed in the lab. Model precision could be increased, at the expense of recall, by increasing the score cut-off threshold.

## Conclusions

The recombinant production of proteins can require multiple experimental rounds of trial and error. To improve the efficiency of such experiments, we have developed RP3Net, an AI model of heterologous protein expression in *E. coli*. RP3Net predicts the results of protein expression as a binary outcome. It was built using the latest foundational models and was trained using a combination of internal experimental results from small-scale AZ expression screens, and publicly available data from the SGC. Using an STP aggregation layer and MLC with large scale purification data enables RP3Net to achieve state-of-the-art performance both on the take-out data from SGC Stockholm and AZ. RP3Net has been experimentally validated on a manually selected set of constructs for viable human drug targets and outperformed third party predictors on that set as well. Ablation studies show that there is no single method that achieves a large performance increase, but rather many small incremental improvements.

This work also underscores the need for large and well curated datasets of soluble protein expression and for the scientific community to agree on how the data should be captured following the FAIR^23,70^ principles, and to establish a protein production ontology. Unfortunately, in the field of protein production there is not yet an equivalent of the PDB for structural biology. Significant time in this project was spent on data curation.

The modelling results may also be further improved by making the model more aware of the experimental conditions, such as *E. coli* host strain, induction methods, time and temperature at which various experimental stages were performed, buffer formulations, etc. This information is largely missing from the currently available data sets.

RP3Net is already deployed and used by the protein scientists at AZ. This publication and the accompanying code repository at GitHub^1^ make the model available to the wider research community, both in industry and in academia.

## Acknowledgements

We would like to thank Susanne Gräslund and Opher Gileadi from SGC Stockholm, and Matthieu Schapira and Peter Loppnau from SGC Toronto for sharing their respective datasets and helping with the curation. We would like to acknowledge colleagues from the Quantitative Biology and Protein Science departments at AstraZeneca for constructive discussions during this project, and David Öling from BioPharmaceuticals R&D at AZ for overseeing the cloning of the experimental constructs. We would like to acknowledge Matthew Hall from the Industry Partnerships team at EMBL-EBI and Birgit Kerber and colleagues from EMBLEM for helping to organise the collaboration; and the EMBL-EBI IT team for maintaining the computational facilities used to train the models. We also acknowledge the funding from the Member States of the European Molecular Biology Laboratory (ARL).

## Methods

### The dataset

Protein production results from AZ, SGC Stockholm and SGC Toronto were used for training and evaluation of the models. AZ and SGC Stockholm report the results of small-scale protein expression testing, after one purification step. In the AZ dataset the outcome is reported as an expression yield category, manually estimated by the scientist who has expressed the protein. In addition, for a subset of constructs, an estimate of the absolute concentration value in mg/L is provided. The estimate is obtained by comparing the size and intensity of the band on the SDS-PAGE gel for the protein of interest with the band for the reference protein of known concentration. This comparison is performed by internal image analysis software. The amino acid sequence of the construct includes affinity and solubility tags; DNA sequences are available for a subset of constructs.

The dataset from SGC Stockholm contains genetic sequences, with tags, annotated with categorical outcomes.

For the bulk of the SGC Toronto data, the outcome is reported as a pipeline position, similarly to PSI/TargetTrack. Importantly, there is no dedicated stage for expression screening: “cloned” is immediately followed by “purified”. Although it can generally be assumed that a protein has to be expressed before it can be purified, it is sometimes the case that producing at larger scale (expression volume) can rescue a construct that failed to yield soluble protein at small scale. For a small subset of SGC Toronto data, small-scale expression screening outcome is also provided as a categorical variable, similarly to SGC Stockholm. Genetic sequences are available for a subset of observations, and tags are included in the constructs.

Graphical overview of the datasets is shown in supplementary figure 2. There are a total of 67,055 unique sequences, covering 5,712 target proteins. Publicly available datasets are significantly larger that the internal AZ data set, SGC Toronto being the largest. Datasets vary in terms of number of constructs per target, availability of genetic sequences vs protein sequences and imbalance between positive and negative outcomes.

To normalise the data across multiple sources and to compute the outcome imbalance, the labels were converted to binary form, with “True” indicating successful production, and “False” failed production. For AZ this binary outcome was computed based on the existing category annotation, estimate of the absolute concentration and manual re-annotation. For SGC Stockholm the binary outcome was derived directly from the existing category annotation. For SGC Toronto, it was derived from the pipeline position.

Historical AZ small-scale expression screening results were reported as concentration range estimates: “0 to 1”, “1 to 10”, “10 to 20”, “20 to 50”, and “above 50” mg/L. This data was converted to binary outcomes as follows. Results in “0 to 1 mg/L” concentration range were annotated as False (not produced). Results within “10 to 20”, “20 to 50”, and “above 50” mg/L were annotated as True (produced). Results in the “1 to 10” range were handled in a special manner. The experiments that had an estimate of absolute concentration were annotated as either True or False by comparing this value with the threshold of 3.5 mg/L. The experiments where the absolute value had not been estimated were re-annotated manually, by re-examining the captured image of the SDS-PAGE gel. AZ data that were collected after April 2023 do not contain manual estimates of the concentration range. Instead, each experiment outcome is manually classified by the scientist into three categories, based on the SDS-PAGE gel: “Passed”, “Not passed” and “Ambiguous”. Outcomes belonging to the “Passed” category were annotated as True, and those belonging to “Not passed” and “Ambiguous” categories as False.

SGC Stockholm reports small-scale soluble expression screening outcomes with manual numeric qualitative annotations by the lab scientist: 0 – no soluble expression, 1 – low soluble expression, 2 – medium, 3 – high and 4 – very high soluble expression. Outcomes from categories 0 and 1 were annotated as False, and with categories 2 and above – as True. Small-scale soluble expression results for SGC Toronto were annotated in the identical manner.

For the SGC Toronto data with pipeline position outcome, results marked with “cloned” were annotated as False, and those with “purified” and beyond - as True.

### Cross validation

To avoid bias towards any particular protein sequence motifs, five-fold cross validation was performed. All constructs were clustered using MMseqs2 v25688290^73^. The affinity and solubility tags were removed from the constructs, and the remaining “target” sequences were clustered. Sequence clusters that contain the AZ results recorded after 1^st^ of September 2023, as well as the constructs used for the experimental validation, were grouped together to form the test set. The remaining constructs were divided into five cross validation subsets, such that each cluster is entirely contained within a single subset. Five-fold leave-one-out cross validation was performed on Model A with SGC Stockholm data, and the worst performing data split was chosen for reporting and for subsequent model development.

### The baseline model

A gradient-boosted decision tree (XGBoost v2.1.3^59^) was used as a baseline model. The input features for the tree were generated by analysing the sequences with ProtParam, as well as predicting global protein properties with Schrodinger API v2021-2^74^, DisEMBL v2.0^75^ and RaptorX^76^. The full list of features and methods used to compute them is given in supplementary table 1. For tools that take protein sequence alignments as input, those were built against the Uniprot database downloaded in February 2016, clustered at 20% cutoff (uniprot20_2016_02^77^).

### Aggregation

Two types of aggregation layer were tested in this work: mean pooling and Set Transformer Pooling (STP^66,67^). For notation, assume that for a sequence of length *N*, the output of the foundational model for each residue *i* is represented with a column vector 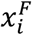 from a *d*-dimensional space, 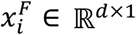. The matrix representation for the entire protein, *X*^*F*^, is obtained by stacking these residue representations along the sequence dimension: 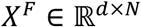. In this notation, mean pooling, which is just averaging all these residue representations, can be written as

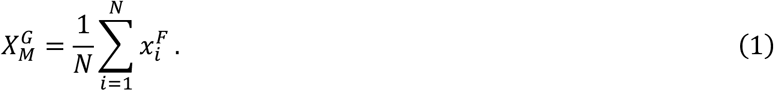

The advantage of mean pooling is that it is simple to interpret and fast to compute. The disadvantage is that it does not have any trainable parameters, or weights, so all training must happen upstream, in the foundational model, or downstream, in the classification head. STP is an example of an aggregation layer with trainable weights. Here, the global representation is the result of performing Multiheaded Attention (MHA^41^) with a seed vector, 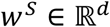, as a query, and residue representations *X*^*F*^as keys and values. Thus,

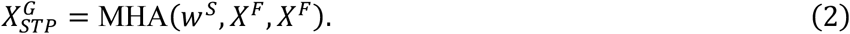

### Multi-headed attention

Computing the MHA^41^ involves weight matrices 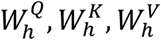 and *W*^O^ for queries, keys, values and outputs, respectively, where *h* = 1.. *H*, and *H* is the number of heads:

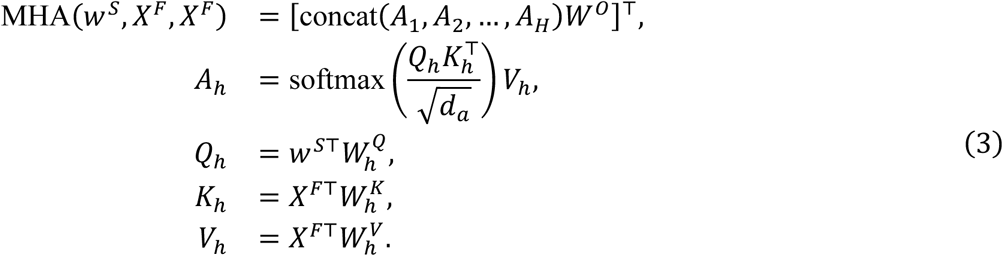

Here, the matrices 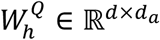 are used to project *w*^*S*^, into *d*_*a*_-dimensional space, 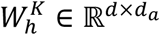 and *W*^*V*^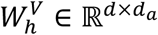 – to project *X*^*F*^into *d*_*a*_-dimensional space, and the matrix 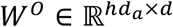 – to project the concatenated single head attention outputs back to the *d*-dimensional space. These matrices, as well as the seed vector *w*^*S*^, are updated during training. The number of heads, *H*, as well as the inputs and outputs dimension, *d*, and the attention dimension, *d*_*a*_, are hyperparameters, that are chosen to maximise model performance on the validation dataset. Matrix transposition, denoted by T, is required to keep the inputs and outputs in column form.

Multi-headed attention is used for the STP aggregation layer in this work, as outlined above. It is also an important part of the transformer architecture, that underpins most of the foundation models.

### RP3Net implementation and training

RP3Net was implemented with PyTorch^78^. Foundation models were downloaded from HuggingFace^79^. Training loop was implemented with PyTorch Lightning^80^. For models C and D, when the foundation model weights were fine-tuned during training, low-rank adaptation (LoRA^81,82^) was used. Early stopping criterion was used, where training is terminated if AUROC for the validation dataset does not improve for 10 epochs. Exact revisions of software packages and foundation models, as well as training run configurations with hyperparameter values, are available in the RP3Net GitHub Repo.

### Meta label correction with purification data

The Meta Label Correction (MLC ^68,69^) framework utilises a larger, noisy, poor-quality dataset to augment the training process of the model that would normally use only a smaller, clean, high-quality dataset. A separate, “teacher” model is trained to predict the corrected soft label from the noisy data and labels. These corrected labels, along with the clean inputs and labels, are used to train the original model, which in this setup is referred to as the “student” model (**Fig. 2B** in the main text).

Formally, we can denote the clean dataset as *D* ≡ {*X*, *y*}, where *X* is the input and *y* is the label. The noisy dataset can be denoted as 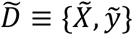, and the corrected labels – as *y*^c^. The student model that predicts the probability of the clean label *y*, based on the clean input *X*, is denoted as *p*_w_(*X*): *P*(*y*|*X*) ∼ *p*_w_(*X*), where *w* are the trainable parameters. In the normal deep learning framework this model is trained by minimising the loss function 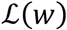 between the true labels and the predicted labels over the clean dataset:

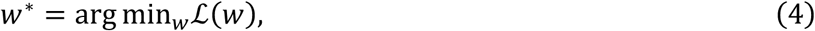

For binary labels that take values of 0 and 1, and cross-entropy (CE) loss, we have

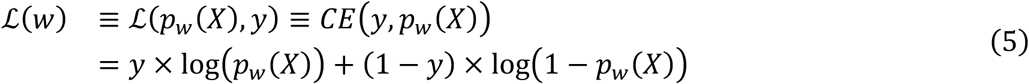

Simple transfer learning would work by substituting *X̃* and *ỹ* and for *X* and *y*, respectively, in equations (4) and (5). Instead, in the MLC framework, the noisy labels *y* are replaced by the corrected labels *y*^c^, modelled by the teacher model, based on the noisy sequences and the noisy labels: 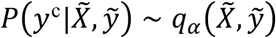, with parameters *⍺*. The loss function 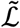 between the corrected labels and the noisy input is obtained by substituting the teacher model in place of *y* in the equation (5):

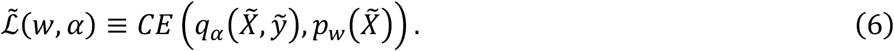

The optimal parameters of the student model *w*^∗^ now depend on the parameters of the teacher model *⍺*:

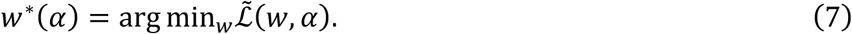

The optimal value *⍺*^∗^ needs to be determined, such that the corrected labels *y*^*c*^ are indeed meaningful in the context of the student model, or, in other words, that the student model trained on the noisy data with corrected labels performs well on the clean data. This can be done by substituting the optimal value of *w* defined by equation (7) in the equation (4):

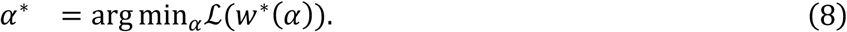

Equations (7) and (8) form the bi-level optimisation problem, that jointly determines the parameters of the teacher and the student models.

In the context of this work, the clean data is a union of the AZ and SGC Stockholm data sets, and the noisy data is SGC Toronto with pipeline position labels. The noisy dataset is thus several times larger than the clean one. On each step of the algorithm several gradient steps through the noisy data (Eqn. 7) are followed by a single step through the clean data (Eqn. 8). The number of noisy steps per single clean step is a hyperparameter. Putting it all together, we get **Algorithm 1** for computing *w*^∗^and *⍺*^∗^.

The teacher parameters *⍺* at step *t* are updated by computing the gradient 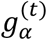 of the clean loss ℒ with respect to (w.r.t) *⍺*. This gradient can be approximated by a formula involving the gradient of the clean loss w.r.t student parameters *w* at step 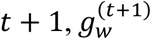, and the matrices of second derivatives (Hessian matrices) of the noisy loss w.r.t *w* and *⍺* at previous steps, 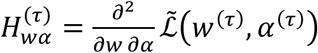:

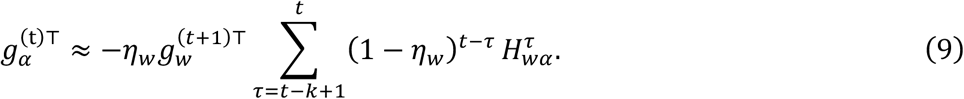

This assumes that gradients are represented as column vectors. For the special case of *k* = 1, the

#### Algorithm 1 Bi-level optimisation of teacher and student model parameters via stochastic gradient descent

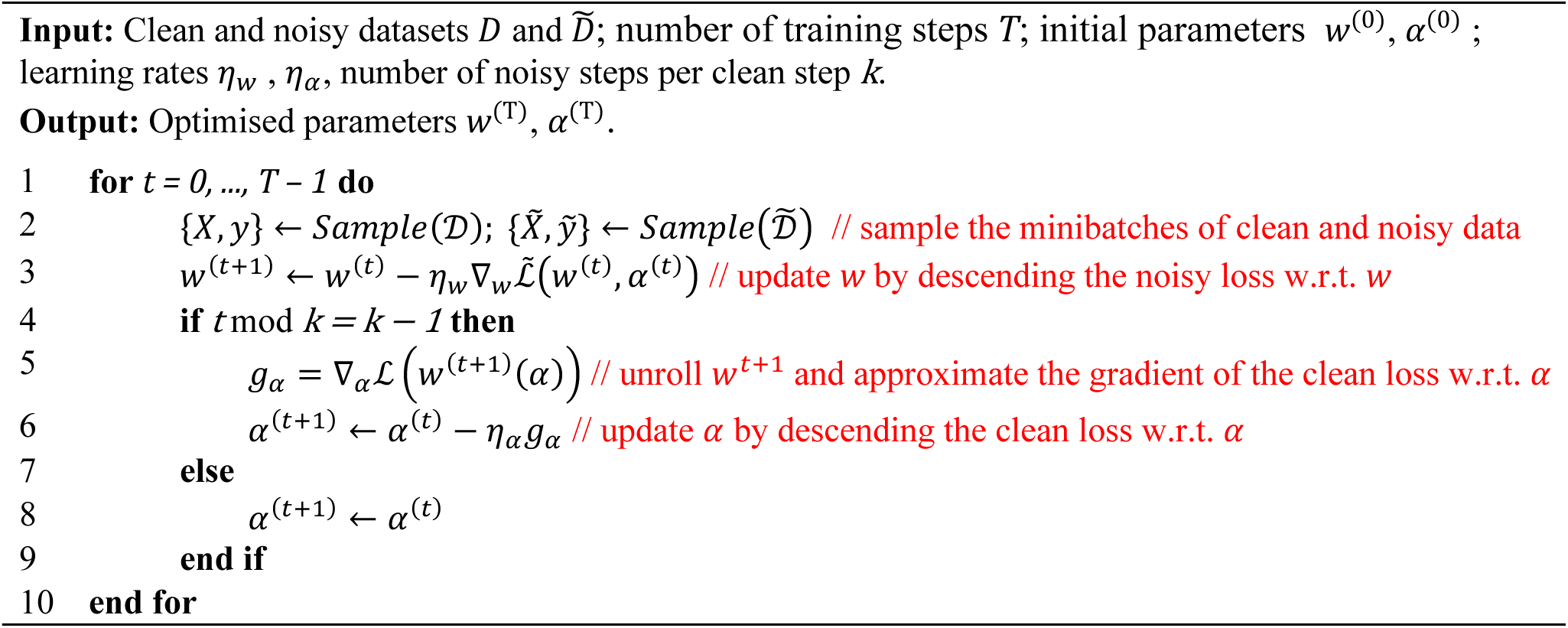

sum in equation (9) is reduced to just 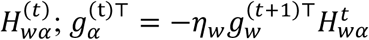.

The CE loss allows for efficient computation of the Hessian *H*_w*⍺*_, by expressing it point-wise as a product of Jacobians, and averaging over the minibatch:

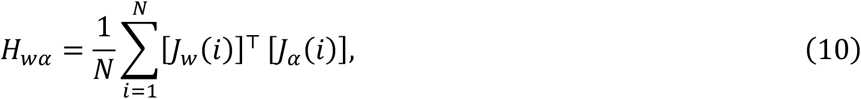

Here, *J*_w_(*i*) is the Jacobian (matrix of derivatives) of the student loss w.r.t *w*, and *J*_*⍺*_(*i*) – the Jacobian of the teacher loss w.r.t *⍺* at input *i*, and *N* is the size of the minibatch.

### Target selection for experimental validation

The target set for experimental validation of the model was curated to include viable human drug targets and exclude proteins that are well known from literature to be successfully expressed. We made sure that neither the protein itself, nor its close homologs, have been deposited in the PDB^52,53^. We have also excluded the target from the validation set if it was referenced from ChEMBL^83^.

OpenTargets^84^ was used to check for viability of a drug target. Twenty thousand human proteins from UniProt^34^ were narrowed down to 454 viable targets. These targets were further curated manually, to have distribution across different target classes and avoid too many DNA-binding proteins. In the end, 46 targets were selected for experimental validation.

Two full length constructs were created per target: one with a TEV-cleavable 6His tag and a GS linker at the N-terminal (MHHHHHHENLYFQGS…), and another one with a GS linker and a 6His tag at the C-terminal (`…GSHHHHHH`). Soluble production of the full-length constructs was predicted with RP3Net. For the targets where both full-length constructs were predicted to fail to be produced, trimmed constructs were generated by iteratively removing residues from N- and C-termini, with a minimum construct length of 50. Trimmed constructs that were predicted to express successfully were included in the experimental validation set. 70% of the set comprised constructs that were predicted to be produced, with the remaining 30% as negative controls. A total of 97 constructs were available for expression testing after taking into account cost constraints and cloning errors.

### Experimental procedures for construct expression

All sequences were codon optimised for *E. coli* and synthesized as synthetic genes (Life Technologies Europe BV) and cloned into backbone vector pET24a. One construct failed during the cloning process. Amino acid and nucleotide sequences of synthesized constructs are found in supplementary table x.

For small-scale soluble expression screening, the plasmid DNA was transformed into competent phage resistant *E. coli* BL21(DE3) cells (New England Biolabs #C2527H) in 96-well PCR plates. The transformation mix was used to directly inoculate 3mL LB media supplemented with 100ug/mL kanamycin in 24 deep-well plates and left shaking at 37°C overnight. Protein expression was auto induced in rich ZYP-8012 media supplemented with 100ug/mL kanamycin in 24 deep-well plates, by inoculating 3mL with 50uL pre-culture and left shaking for 3 h at 37°C followed by 24 h at 18°C. After harvest (4000xg, 5min, 4°C), the pellets were lysed with 900uL lysis buffer (40mM HEPES, 300mM NaCl, 5mM imidazole, 10% glycerol, 1mM TCEP, 0.1% DDM, 0.2mg/mL lysozyme, DNAse & protease inhibitors) and freeze-thawed once. The lysate was cleared by centrifugation (4000xg, 30min, 4°C) before subjecting to a one-step Nickel affinity purification using an automated bead-based platform. The protein was captured on the magnetic beads for 30min at 4°C, followed by two wash steps to wash off unbound proteins (40mM HEPES, 300mM NaCl, 5mM imidazole, 10% glycerol, 1mM TCEP) and eluted in 100uL elution buffer (40mM HEPES, 300mM NaCl, 300mM imidazole, 10% glycerol, 1mM TCEP). 10uL of the elution was loaded onto Nu-PAGE Bis-Tris gels (Invitrogen) together with Novex Pre-stained protein marker (Invitrogen) and 5ug of an internal standard protein. The gels were stained in Der Blaue Jonas (GRP) and analysed using the densitometry software Image Lab (BioRad). The protein yield (mg/L) was estimated from the relative quantity. The “Passed”, “Not Passed” and “Ambiguous” outcome annotations were provided manually by the lab scientist, based on the relative thickness and brightness of gel bands. Annotations from two separate biological replicates are shown in supplementary Table X. Gels from one of the two experiments are shown in Supplementary Figure X. The maximum yield from the two experimental runs was used as the ground truth for the model evaluation.

## Footnotes

1 www.github.com/RP3Net/RP3Net

