## Supplementary material for "RP3Net: a deep learning model for predicting recombinant protein production in *Escherichia coli*": sup_fig_1_expression_gels.pdf

### Results: SDS- PAGE gels

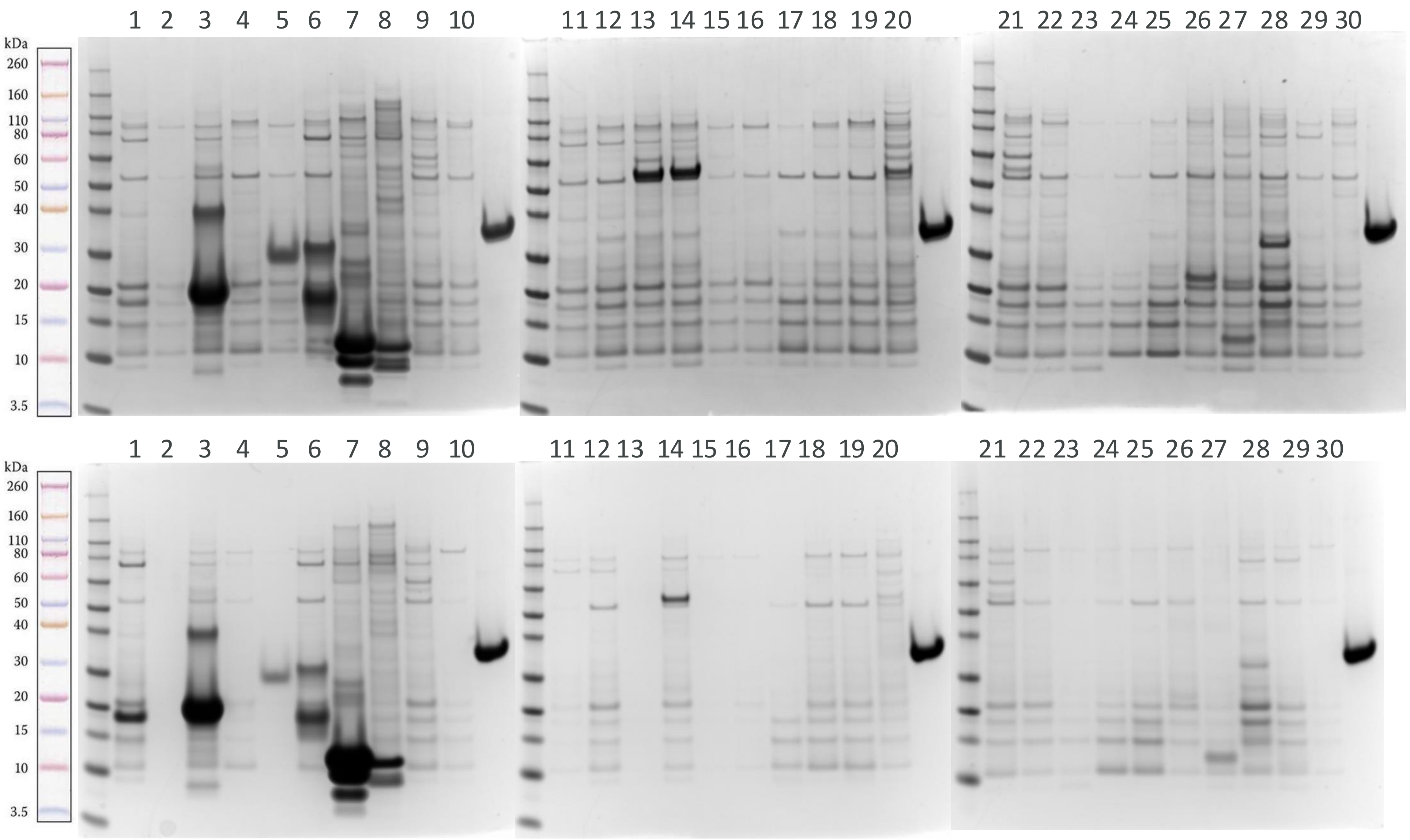

### Results: SDS- PAGE gels

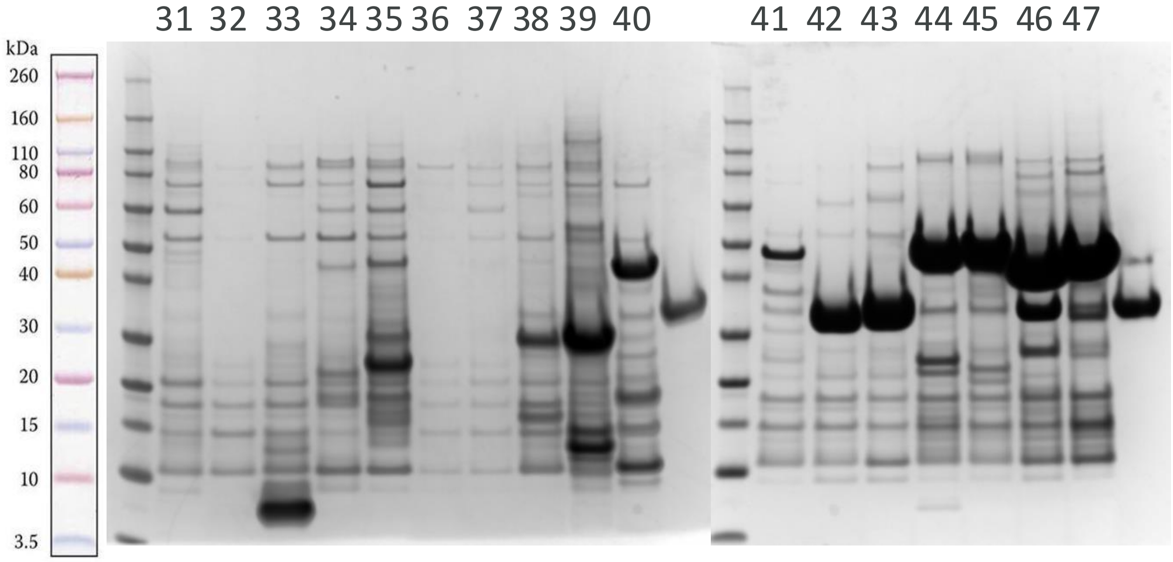


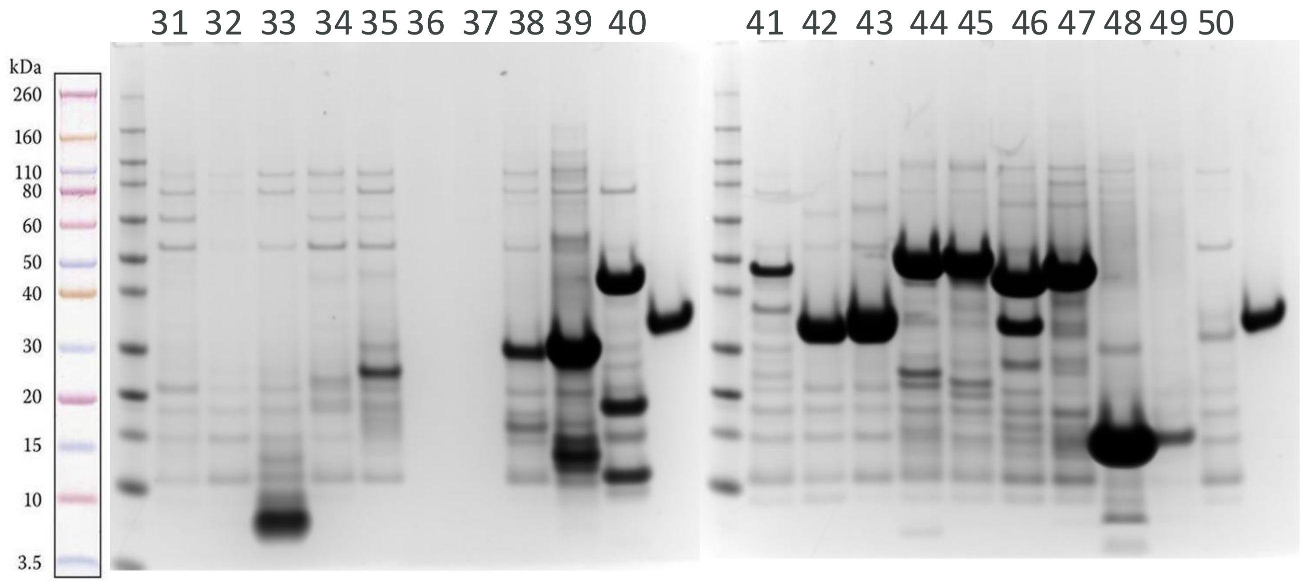

### Results: SDS- PAGE gels contd.

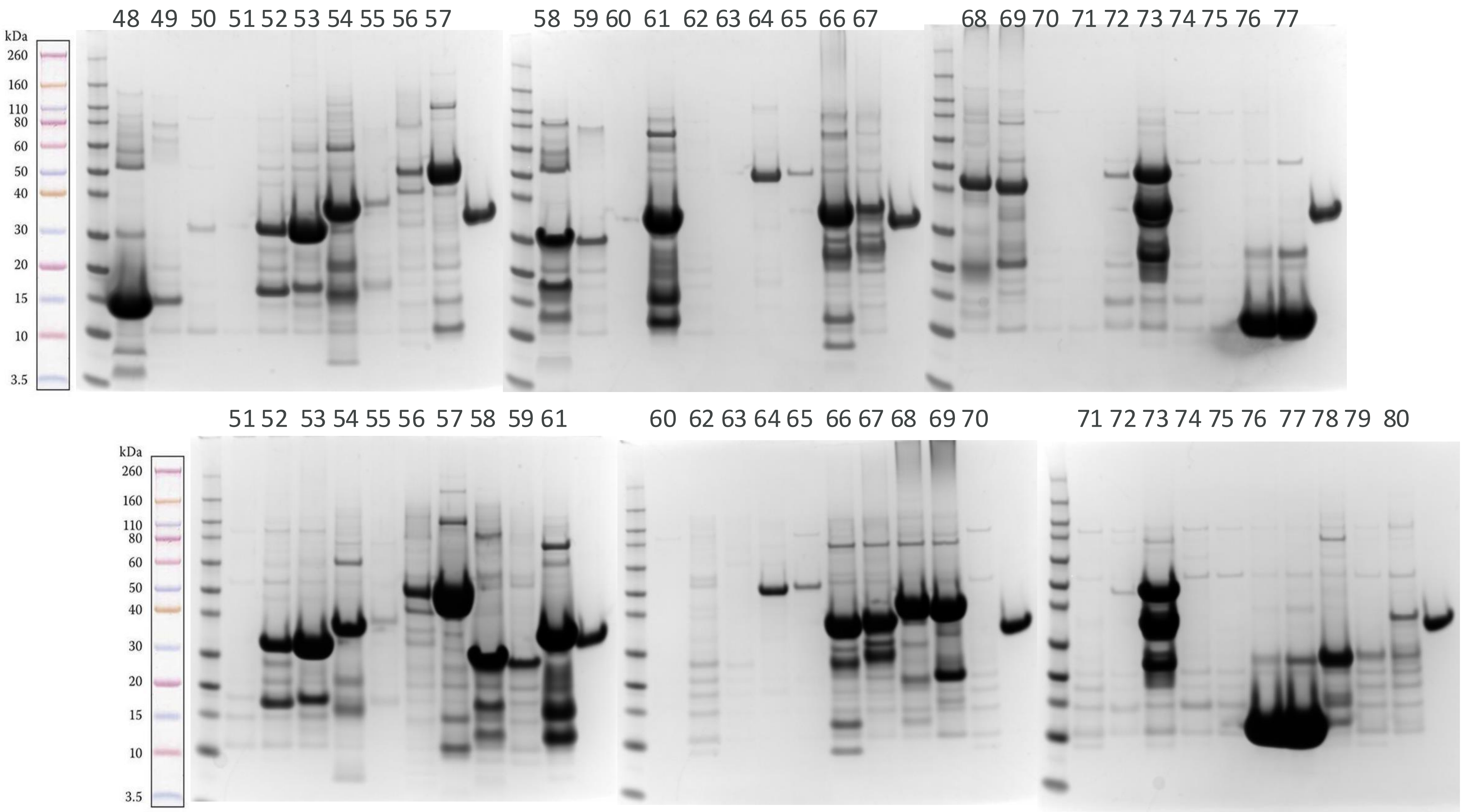

### Results: SDS- PAGE gels contd.

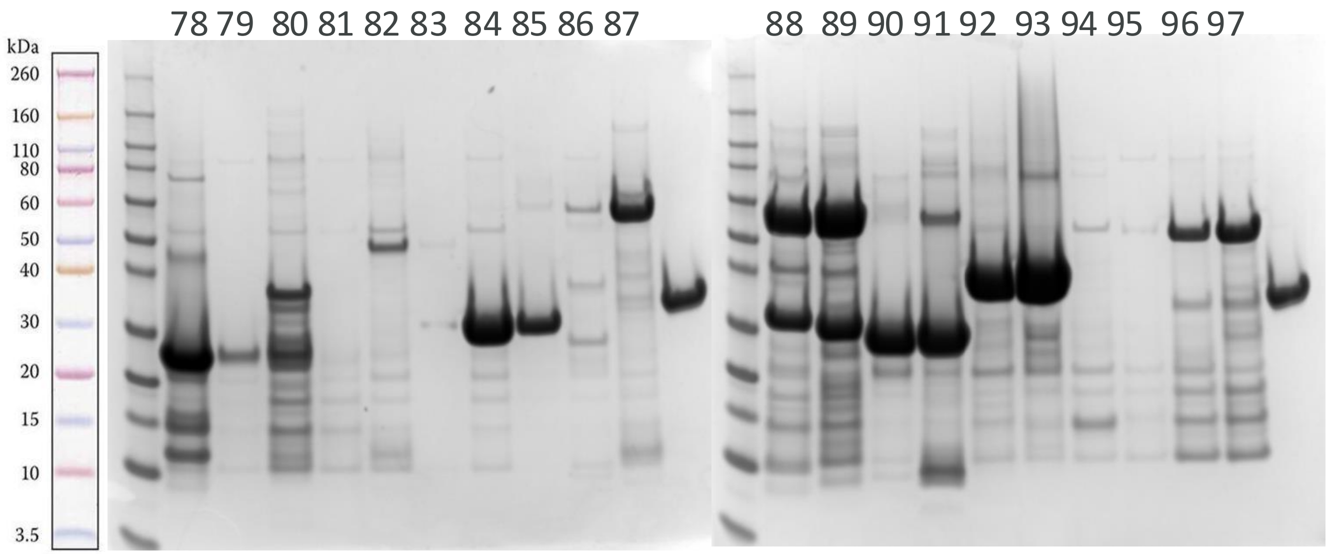


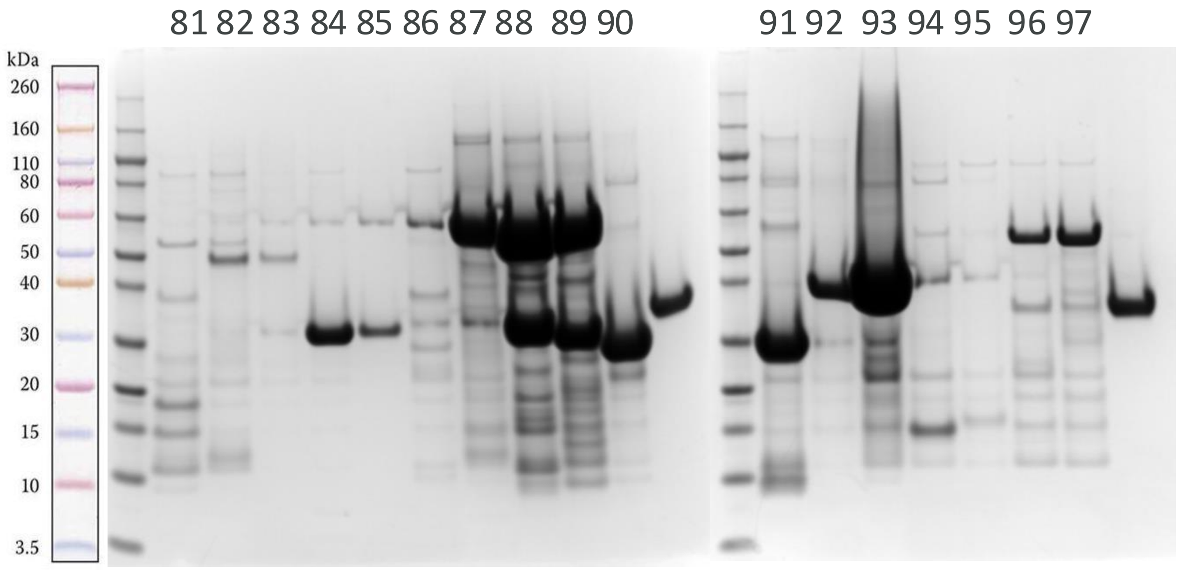
