## Supplementary figures and images for "RP3Net: a deep learning model for predicting recombinant protein production in *Escherichia coli*"

### supp_fig_2_ds_overview.png

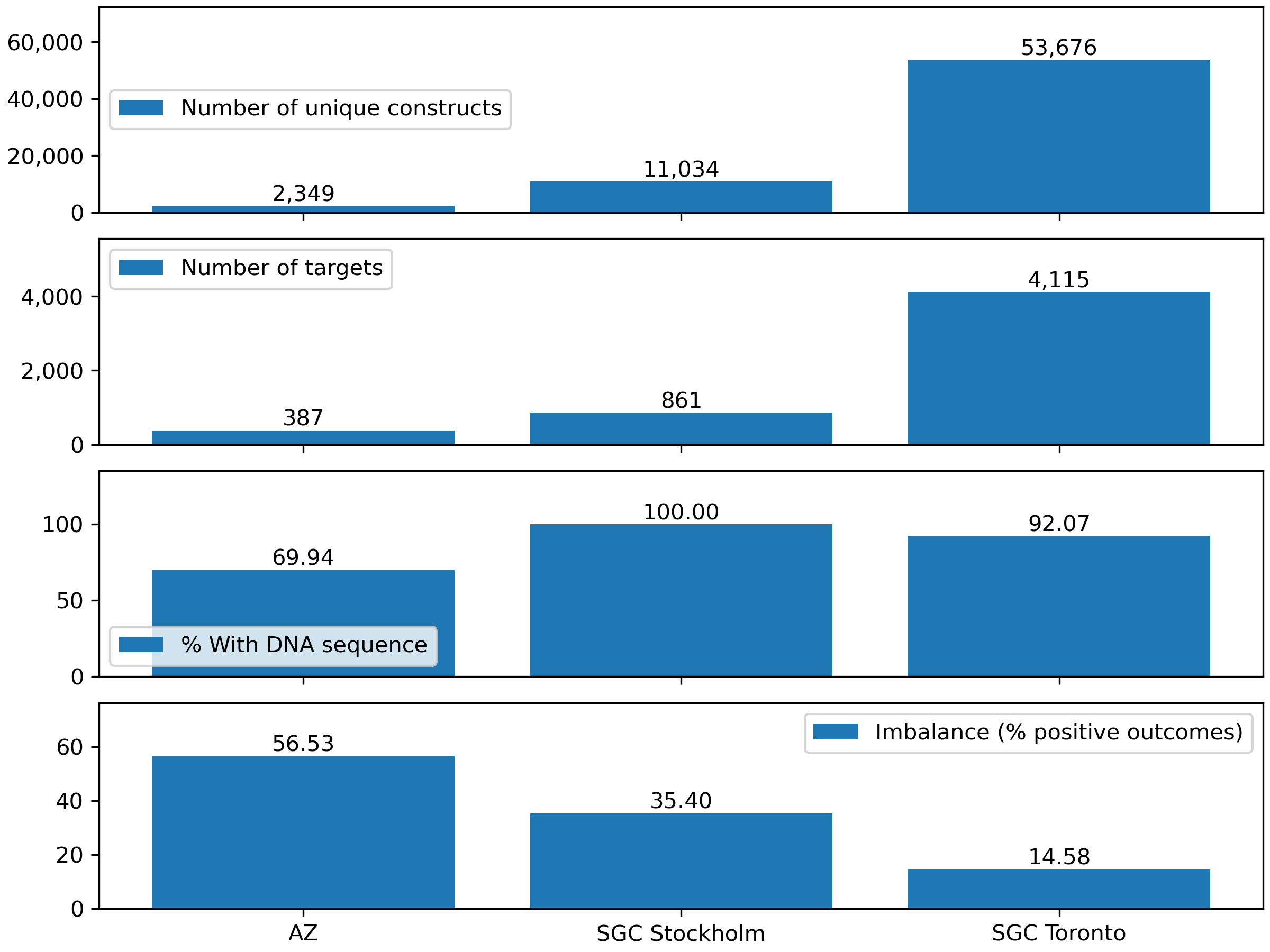
